# Accelerated biomass loss in western US forestlands due to shifting fire regime

**DOI:** 10.1101/2025.08.22.671876

**Authors:** Shike Zhang, Neil Carter, Joseph O. Sexton, Jonathan A. Wang, Panshi Wang, Kai Zhu

## Abstract

Wildfires influence the distribution of biomass across the Earth’s surface and drive losses of carbon from the land surface to the atmosphere. Although the global budget between terrestrial and atmospheric carbon pools is comparatively well understood (Jones et al., 2024; MacCarthy et al., 2024), the growing size and severity of wildfires present an increasing challenge to regional carbon accounting. To quantify vegetation biomass across arid systems in the western US, we used remote sensing data to estimate the aboveground live biomass density (Mg ha^-1^) at 30-meter resolution across the states of Utah and Nevada annually from 2000 to 2022. Time series biomass maps showed accelerated loss of terrestrial carbon to the atmosphere as a result of increasing wildfire, with annual biomass burnt increasing at 0.105 ± 0.024 Mt yr^-1^ (mean ± standard deviation) after 2015. Recent shifts in the wildfire regime show a transition from predominantly early-season fires in low-biomass grasses and shrubs to late-season fires in higher-biomass forestlands. The proportion of total biomass loss attributed to wildfires in forestland areas increased from an average of 76% per year before 2015 to 94% in the years that followed. Furthermore, we found that recent droughts contributed to increased biomass loss in forestland areas, whereas biomass in non-forestland areas appeared unaffected by drought conditions. These findings underscore the escalating impact of climate change on fire regimes, highlighting the urgent need for adaptive land management strategies to mitigate carbon loss and preserve ecosystem resilience in the face of ongoing aridification.

## 1. Introduction

Wildfires play a critical role in developing and maintaining global ecosystems, significantly influencing their structure and function (Bowd et al., 2021). The prevalence of wildfires across different terrestrial landscapes, ranging from forests to shrublands and grasslands, results in significant biomass loss (Foster et al., 2020). This loss not only impacts ecosystem structure, but also contributes to atmospheric carbon emissions, further strengthening feedback between wildfire activity and global climate dynamics (Landry & Matthews, 2016). However, despite the widespread recognition of wildfire impacts (van Wees et al., 2022; Wang et al., 2021), there remains a critical knowledge gap in quantifying the direct biomass loss caused by wildfires under climate change in different land covers. While extensive literature exists on wildfire emissions (Y. Liu & Ding, 2024; van der Werf et al., 2017; van Wees et al., 2022; Zhao et al., 2023), these studies typically rely on emission factors rather than direct measurements of biomass change. A fuller appraisal of biomass loss and its mechanistic drivers are needed to understand future ecosystem dynamics under change, and to anticipate the consequences of those changes on ecosystem functioning and human wellbeing.

Growing evidence suggests that climate change is driving increasing wildfire activities (Senande-Rivera et al., 2022) in increasingly arid regions (Grillakis et al., 2022; Mahmoud et al., 2023). Climate change might push critical parts of the ecosystem to tipping points (Lenton et al., 2024), which in turn could change wildfire regimes (Newberry et al., 2020), including biomass loss. Globally, wildfires have been shown as a major contributor to carbon emissions (Jones et al., 2024; MacCarthy et al., 2024). Wildfires release large amounts of stored carbon from trees, shrubs, and other plant materials, contributing up to one-third of global carbon emissions (Y. Liu et al., 2014). This sudden release not only elevates atmospheric CO₂ levels, intensifying climate change (Solomon et al., 2009), but also weakens ecosystems’ long-term carbon sequestration capacity (Wang et al., 2021).

Biomass-dense regions, such as forests, play a key role in this process. For example, forests are widely regarded as a natural climate solution due to their ability to absorb CO_2_ through photosynthesis and function as carbon sinks (Pan et al., 2024). However, when wildfires occur, these forest carbon sinks can rapidly shift into carbon sources, which is often acknowledged but rarely quantified (Anderegg et al., 2022; Landry & Matthews, 2016; Loehman, 2020). The growing extent and intensity of wildfires heighten the risk of a self-reinforcing feedback loop, in which increased carbon emissions lead to further warming, fueling even more frequent and severe fires—a cycle that accelerates global climate change (Mehmood et al., 2020). Given projected climatic trends, this cycle poses a significant threat to forest carbon stocks and their ability to mitigate climate change through carbon sequestration (Liang et al., 2017). Understanding the impact of climate change-driven wildfires on biomass is essential for accurately assessing carbon dynamics and enhancing ecosystem resilience.

Broader land cover dynamics, influenced by increased wildfire frequency and global environmental changes, necessitate comprehensive research across diverse ecosystems. Intact forests are particularly important to this accounting due to their role in the global carbon cycle and the ecosystem services they provide (Pan et al., 2024), with their carbon storage capacity threatened by increasing wildfire frequency and intensity (Jain et al., 2024; Wang et al., 2024). Focusing on wildfires in forest areas is crucial because densely packed, dry underbrush can lead to more severe fires compared to environments with moist or sparse underbrush or grasslands (Jolly et al., 2014; Loudermilk et al., 2022; Varner et al., 2015). Yet, while forests are critical, they only comprise about a third of all land, and other land cover types play significant roles in the ignition and suppression of wildfires, necessitating comprehensive research across diverse land cover types (Cochrane & Ryan, 2009; Mugabowindekwe et al., 2023; Tucker et al., 2023). Recently, more frequent wildfires have caused long-term land cover transformation from forests to other land cover types in the western US (Coop et al., 2020; Hessburg et al., 2016). This change is largely attributed to the complex interplay of various global change factors, including climate change (Foster et al., 2020; Parks et al., 2016), land use change (Radeloff et al., 2018), and human demographic shifts (Keeley & Syphard, 2020). The land cover transformation resulting from these changes calls for studies expanding beyond forest areas to address the dynamics in other land cover types.

To address this call, we examine how wildfires affect biomass and land cover across different land cover types in Utah and Nevada in the western US. We focus on this region because it has diverse land cover types that are also highly vulnerable to severe wildfires. Both states rank among the fastest-growing in the nation (Bureau, 2020), leading to rapid urbanization and expansion into the wildland-urban interface (Radeloff et al., 2018). As human development expands into previously remote areas, the likelihood of wildfire ignitions from human activities increases (Balch et al., 2017; Calkin et al., 2014; Keeley & Syphard, 2020; Westerling et al., 2011). This expansion exposes ecosystems that were once less affected by human presence to a growing risk of wildfire. These wildfires threaten both natural ecosystems (Jakus et al., 2017) and human communities (Mallia et al., 2015), leading to biomass burning that elevates carbon emissions and degrades air quality (Butt et al., 2020), with significant consequences for human health (Johnston et al., 2021). At the same time, increasing aridity (Abatzoglou & Williams, 2016; Williams, Cook, et al., 2022) in these two states exacerbates wildfire risk (Barea-Azcón et al., 2023; Khatri et al., 2018). Given the diverse land cover types in Utah and Nevada, the rapidly expanding wildland-urban interface, and the widespread cheatgrass invasion (Smith et al., 2023), these two states provide an ideal setting to study the complex regional impacts of wildfires across different landscapes. Climate change has led to an increase in the area burned by wildfires in forestland ecosystems across the western US (Abatzoglou & Williams, 2016), with most research concentrating on these expanding burn areas (Magerl et al., 2023; Urbanski et al., 2018; Xu et al., 2022), while the carbon consequences of these wildfires still need further investigation.

We have two main hypotheses that guide this work. First, we hypothesize that biomass loss during wildfires has increased in recent years due to more frequent and severe wildfires, particularly driven by climate-induced aridification and prolonged droughts. Second, we hypothesize that forests have lost more total biomass than shrublands and grasslands, despite the latter often experiencing larger burn areas. To test these hypotheses, we leverage high-resolution (30 m), annual (2000-2022) remote sensing data to quantify biomass loss by integrating wildfire occurrence data with biomass estimates and identify trends across major land cover types such as forests, shrublands, and grasslands. We further assess how differences in vegetation structure and fuel availability influence biomass burning and fire regime shifts, exploring both interannual variability and long-term trends. By examining these patterns, our study aims to improve the understanding of wildfire impacts on biomass dynamics and fire regimes, contributing to carbon management and ecosystem conservation efforts.

## 2. Materials and methods

### 2.1. Study area and data sources

Our study area is located in the western US, encompassing regions of Utah and Nevada. These two arid states cover 212,761 km^2^ and 284,332 km^2^, respectively, with highly variable topographic relief ranging from 147 m to 4,120 m. Since the 2010s, Utah and Nevada have experienced some of the highest population growth rates in the US of 15.0-66.3% per year (Bureau, 2020). This rapid growth is anticipated to expand the wildland-urban interface (Schug et al., 2023), increasing the potential for extreme fire behavior (Birch & Lutz, 2023). Besides the population growth and the wildland-urban interface increase, the invasion of cheatgrass, which leads to large wildfire,s is also a problem in these two states (Smith et al., 2023). Between 1984 and 2020, Utah and Nevada recorded a total of 1,733 wildfires (> 202 ha), averaging 47 fires per year and burning an average of 201,711 ha annually.

To better understand the spatial and temporal patterns of these wildfires, we utilized geospatial data from the Monitoring Trends in Burn Severity (MTBS) program, which provided wildfire boundaries, wildfire occurrence dates, and burn area estimates for wildfires larger than 1,000 acres occurring from 2002 to 2020 (January 1, 2002, to December 31, 2020). Out of 901 wildfires in Nevada and Utah recorded during the study period, 900 were selected for analysis. The remaining one fire was excluded due to MTBS mislabeling (Map ID = 3972). The total burn area analyzed in this study is approximately 4,429,421 ha, which is about 8.7% of the total area of Utah and Nevada. Although there is some debate that MTBS tends to overestimate fire polygons by 16–35% (Kolden et al., 2015), this study used biomass loss to estimate the impact of wildfires, so the inclusion of unburnt areas does not significantly affect the accuracy of our findings. Moreover, the consistent methodology used for fire polygons in MTBS ensured robustness against temporal and spatial variability (Dennison et al., 2014).

The land cover data was obtained from the National Land Cover Database (NLCD) by the U.S. Geological Survey (USGS). The NLCD program provides land cover and land cover change at 30-meter resolution in the US. NLCD uses surface reflectance time series from the Landsat series of satellites as inputs for land cover classification and post-classification processes employing random forest models (Wickham et al., 2023). NLCD includes map products characterizing land cover types in eight years (Homer, 2020) from 2001 to 2019 (2001, 2004, 2006, 2008, 2011, 2013, 2016, and 2019) with a 30-meter resolution. We aggregated the NLCD categories into four major classes: forestland (including woody wetlands, evergreen forest, deciduous forest, mixed forest), shrubland (including shrub/scrub), grassland (including herbaceous, emergent herbaceous wetlands), and others (including developed - open space, developed - low intensity, developed - medium intensity, developed - high intensity, open water, cultivated crops, hay/pasture, barren land, perennial snow/ice). In this study, we overlaid each MTBS fire polygon onto the most recent NLCD map available before the wildfire occurred (excluding the year that the wildfire happened).

The biomass data for Utah and Nevada were obtained from terraPulse, Inc., which provides time series maps of aboveground biomass at a 30-meter spatial resolution with annual coverage from 1984 to 2021. This dataset is validated by field plot data from the United States Forest Service (USFS) Forest Inventory and Analysis (FIA) program (Cao et al., 2025). The terraPulse dataset includes total aboveground live biomass (Mg ha⁻¹) and is derived from a model trained on NASA’s Global Ecosystem Dynamics Investigation (GEDI) L4A footprint level aboveground biomass density and other environmental datasets, such as tree canopy cover, land cover, and disturbance history. The terraPulse model utilizes gradient-boosting regression trees (GBRTs) for prediction and incorporates uncertainty estimation through Monte Carlo sampling and model-based error analysis. To enhance precision, biomass estimates undergo post-processing steps, including water body masking. In this study, we overlaid each MTBS fire polygon onto two terraPulse biomass maps from different years: one from the year before the wildfire occurred and the other from the year after. Each biomass map is accompanied by a corresponding uncertainty map, which has the same spatial structure as the biomass maps, except that each pixel’s value represents the uncertainty associated with the corresponding pixel in the biomass map. We used Monte Carlo simulations to quantify the overall uncertainty in biomass estimates for specific regions in Utah and Nevada (SI §1).

The drought indices were obtained from the Gridded Surface Meteorological (gridMET) dataset (Abatzoglou, 2013). GridMET is a high-resolution (∼4-km) daily meteorological dataset for the continental US from 1979 to the present (Abatzoglou, 2013). gridMET supports a range of drought indices computed on multiple time scales, including 14 days, 30 days, 90 days, 180 days, 270 days, 1 year, 2 years, and 5 years. The drought indices include the standardized precipitation index (SPI), the evaporative drought demand index (EDDI), the standardized precipitation evapotranspiration index (SPEI), the Palmer drought severity index (PDSI), and the Palmer Z Index (Z). (SI §2)

### 2.2. Data preparation

#### 2.2.1. Temporal alignment of wildfires

To assess the impact of wildfires on biomass, we focused on comparisons between pre- and post-fire biomass status. We used remote sensing data from the year before the wildfire as the pre-fire state and data from the year after the wildfire as the post-fire state. We excluded the year the wildfire occurred to ensure consistency in our comparisons. This exclusion was necessary because some wildfires occurred before the year’s map was produced, while others occurred after. Therefore, excluding the year of the wildfire provided a standardized approach for our analysis.

Given the temporal discontinuity of land cover data, this study uses the land cover map from the date closest to the wildfire prior to its occurrence as a proxy for land cover conditions during the event. This is because, in the absence of external disturbances, the land cover remains unchanged (Fig. S1), making it reliable to use land cover maps from a few years before the wildfire as a representation of the land cover at the time of the wildfire.

#### 2.2.2. Spatial alignment of wildfires, land cover, and biomass data

To assess biomass loss due to wildfires, we integrated MTBS fire polygons, annual terraPulse biomass maps, and NLCD land cover maps to create a spatiotemporally aligned dataset BioLandPix, capturing information on biomass (Bio), land cover (Land), and pixel-level spatial resolution (Pix). BioLandPix includes biomass estimates before and after each wildfire, as well as the land cover type before each wildfire, for each pixel from 2002 to 2020 (January 1, 2002, to December 31, 2020) in Utah and Nevada. The integration has three steps: 1) extracting pre- and post-fire biomass based on MTBS, 2) retrieving land cover data for all pixels within each wildfire boundary, and 3) filtering out wildfires, which are mistakenly labeled.

First, we obtained MTBS polygons to define the boundaries of wildfire-affected areas. We overlaid these MTBS polygons onto the terraPulse annual biomass maps to extract biomass estimates within the limits of each wildfire event (Fig. S2). From the biomass data (pre-wildfire only) of each wildfire, we extracted the latitude and longitude coordinates of the geometric center of each pixel within the MTBS polygons. The geometric center served as the representative geographic location for the entire pixel area.

Second, we matched pre-fire biomass pixels to the nearest pre-fire NLCD map to determine land cover types. We extracted geographic coordinates from the terraPulse biomass maps for each wildfire. These coordinates were then mapped onto the NLCD map to identify the land cover type for each pixel in the fire area prior to the wildfire.

Last, to ensure data completeness in BioLandPix, we implemented a pixel-by-pixel filtering process for each wildfire. If any of the pixels lacked pre-fire or post-fire biomass data or land cover data, we marked the pixel as incomplete and excluded it from further analyses. What’s more, we examined a total of 901 wildfires, of which one (Map ID = 3972) was identified as mislabeled by MTBS. Ultimately, BioLandPix included 900 wildfires (99.9 % of the total wildfires), encompassing 6,652,739 ha (73,919,327 pixels), from January 1, 2002, to December 31, 2020.

### 2.3. Geospatial and statistical analyses

#### 2.3.1. Change in interannual wildfire loss

To analyze the temporal trend of biomass loss and to detect if there is a sudden change in biomass loss, we used segmented regression to detect tipping points that indicate significant shifts in biomass dynamics. The number of breakpoints of annual biomass loss in the research area was determined based on exploratory data analysis, which suggested a single, clear change point in the time series. Segmented regression fits a model where the relationship between the response variable and one or more explanatory variables is piecewise linear. It assumes that the data is represented by multiple connected straight lines, known as segments, with breakpoints or changepoints where the slope of the line changes (Muggeo, 2008). This method is best suited for our analysis as it allows us to examine potential threshold-driven shifts in biomass loss, which may arise as climate change pushes ecosystems to tipping points, altering flammability thresholds and abrupt changes in wildfire biomass loss. By applying segmented regression, we can identify the tipping point in biomass loss, which is crucial for understanding when and how wildfire impacts escalate beyond a critical threshold.

The equation for the segmented regression model applied in this study can be expressed as:

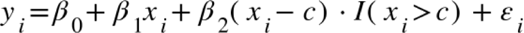

where *β_0_* is the intercept of the first segment, *β_1_* is the slope of the first segment, *β_2_* is the change in the slope after the breakpoint (c), ^*I*(*x*_*i*_ > *c*^) is an indicator function that equals 1 if (*x*_*i*_ > *c*) and 0 otherwise, *ε*_*i*_ is the error term, assumed to be normally distributed: *ε_i_* ~ 𝓝(0, *σ*^2^). Because the model is continuous at (c), the actual slopes for *x*_*i*_ > *c* are (*β_1_*+ *β_2_*), ensuring seamless transition between the two segments. The change in the slope is determined by *β_2_* and the indicator function, which activates only for points beyond (c) (SI §5).

#### 2.3.2. Relationship between aridification and biomass loss during wildfires

To further understand the role of drought in shaping wildfire outcomes, we investigated how pre-fire drought conditions influence wildfire biomass loss across different land cover types. Previous studies suggest drought conditions play an important role in affecting the burn area of wildfires, as they directly impact vegetation moisture levels and fuel availability (Abatzoglou & Williams, 2016; Littell et al., 2016; Mansoor et al., 2022; Williams, Cook, et al., 2022). In this study, we analyzed the relationship between drought conditions and wildfire biomass loss across different land cover types. We extracted drought indices (EDDI, SPI, SPEI, PDSI, and Z) from gridMET at multiple time scales (14 days, 30 days, 90 days, 180 days, 1 year, 2 years, and 5 years) prior to wildfires and at their locations, along with the corresponding land cover types. By using only pre-fire drought data, we ensured that drought was treated as the cause and wildfire biomass loss as the effect.

To systematically explore the relationships between various drought indices and biomass loss, we employed the multivariate information-based inductive causation (MIIC) algorithm, a data-driven approach for causal analysis. The MIIC algorithm infers graphical networks that represent direct and potentially causal relationships among variables based on information-theoretic principles (Verny et al., 2017). By iteratively identifying and removing redundant connections, MIIC reconstructs a concise and interpretable network structure without relying on temporal ordering or experimental interventions. It can estimate conditional mutual information and does not require hyperparameter tuning or pre-imputation for missing data, making it well-suited for ecological datasets that are often incomplete or noisy (Sella et al., 2025). Compared to traditional correlation-based methods, MIIC enables the detection of both direct and indirect dependencies, and under certain conditions, can infer causal directionality (Verny et al., 2017). In this study, we used the MIIC-derived network to identify a subset of drought indices that are most directly associated with biomass loss, while accounting for the complex interdependencies among indices themselves. Rather than treating all indices as independent predictors, MIIC allows us to disentangle direct from indirect relationships, effectively reducing redundancy and simplifying the variable space. After incorporating all the drought indices and biomass loss data into the algorithm, we obtained a graphical network indicating the potential causal relationships between the drought indices and biomass loss. This approach offers a clearer view of which indices carry unique information about biomass loss, and provides a more interpretable structure for ecological analysis.

## 3. Results

### 3.1. Biomass and land cover conditions

Utah and Nevada’s biomass distribution across different land cover types has been remarkably stable over time. The total biomass in Utah and Nevada remained overall unchanged at approximately 540 Mt over the years, experiencing only slight fluctuations (Fig. S3). The biomass stored in different land cover types was also stable. The dominant land cover type in Utah and Nevada is shrubland, occupying approximately 64.8% of the total land area (Fig. 1b). Forestland, although it makes up only about 15.75% of the total land area, has the most biomass, accounting for 47.95% of the total biomass in the region (258.84 Mt total and 32.3 Mg ha⁻¹ on average) (Fig. 1b and 1c). This is slightly more than the shrubland’s 46.14% of the total biomass, approximately 249.04 M and 7.5 Mg ha⁻¹. While for the grassland, it only covers 9.36% of the land area and only accounts for 5.91% of the total biomass stored in this area, which is about 31.9 Mt and 6.7 Mg ha⁻¹. Compared to wildfires in shrubland and grassland, wildfires in forestland areas experienced more severe damage and greater biomass loss (Fig. 1a).

**Fig. 1.**
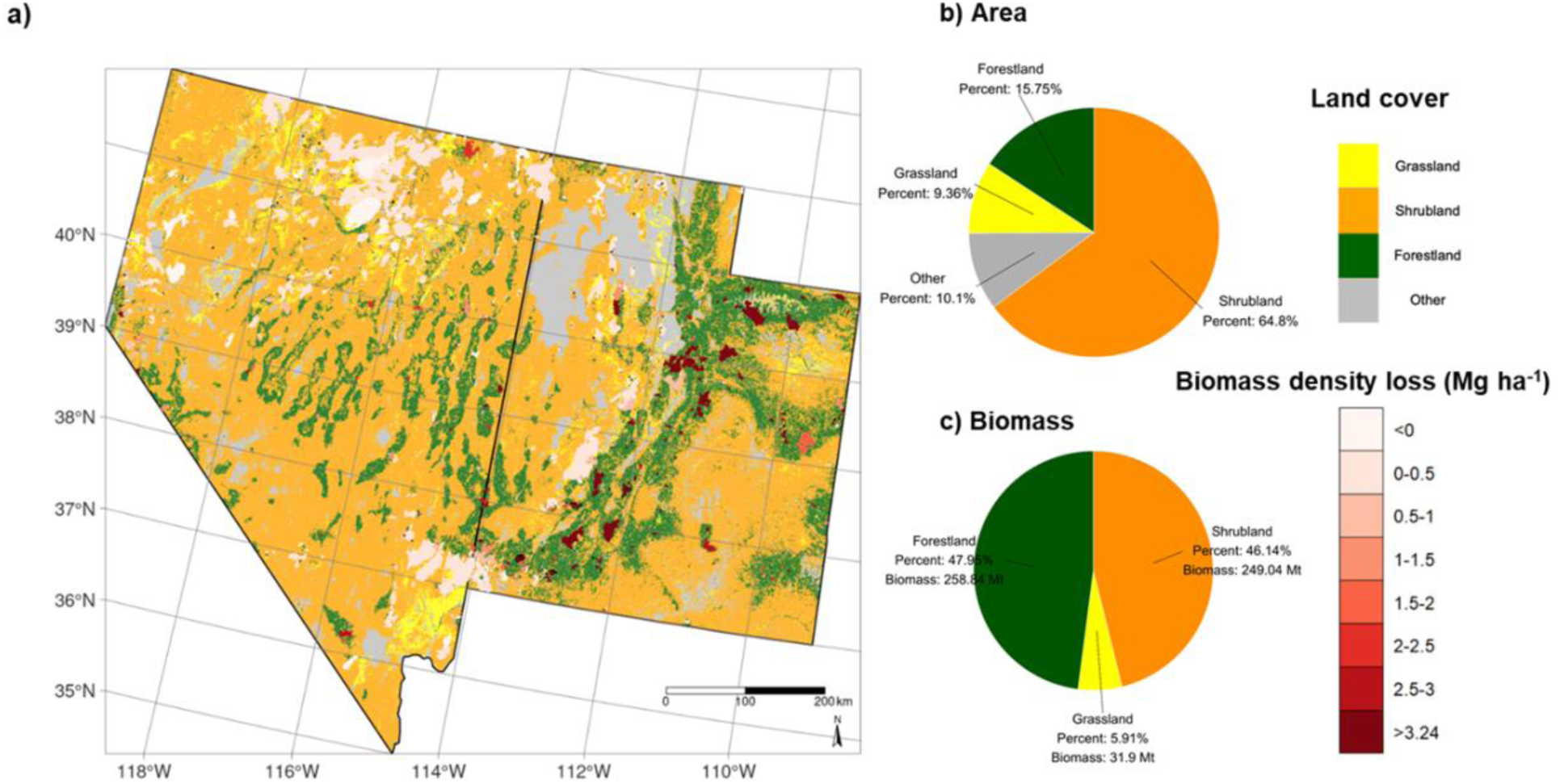
Overview of study area characteristics and biomass loss due to wildfires. **(a)** A map of land cover distribution and biomass density loss caused by wildfires. Land cover shown is based on the 2019 NLCD data. Superimposed on the raster plot are polygons shaded in varying intensities of red, representing wildfire burn areas. The colors indicate each fire’s biomass density loss (Mg ha^-1^), calculated using terraPulse biomass data. The biomass density loss for each polygon is determined by measuring the difference in total biomass before and after each wildfire and dividing this difference by the total burned area provided by MTBS. Accompanying the figure are two pie charts: **(b)** Percentage of each land cover type by area within the study area using the 2019 NLCD land cover data, and **(c)** Percentage of biomass for each land cover type. This latter percentage is derived by overlaying the NLCD land cover maps with terraPulse’s biomass remote sensing data collected over multiple years and averaging the annual total biomass for each land cover type.

### 3.2. Temporal trends and tipping points in biomass burning impact

The segmented regression on annual biomass loss during wildfires reveals a significant tipping point occurring in 2015 (*c*, *p* < 0.01). This presents an important shift in the biomass loss pattern caused by wildfires. Before 2015, the trend of biomass loss caused by wildfires was not significant (*p* > 0.1). After 2015, there was a significant increase in the total biomass loss caused by wildfires in Utah and Nevada (*p* < 0.01), with annual biomass burnt increasing at 0.105 Mt yr^-1^ (*β_1_* + *β_2_*) with a standard deviation of 0.024 Mt yr^-1^. In 2015, the total biomass burnt was recorded at 68,565 ± 7,219 Mg (0.069 ± 0.007 Mt), marking the start of this upward trend (Fig. 2a). Over the following years, this number has steadily increased (*β_1_* + *β_2_* = 0.105 Mt yr^-1^), reflecting the increasing damage of wildfires to the local ecosystem. By the end of the most recent measurement period in 2019, the annual biomass loss had risen substantially to 465,999 ± 17,199 Mg (0.466 ± 0.017 Mt), which is about 6.8 times the biomass loss in 2015. Compared to the previous 12 years, where the average biomass burnt was 137,895 ± 3,214 Mg yr^-1^, the period from 2015 to 2019 experienced a much higher average biomass loss of 344,310 ± 6,608 Mg yr^-1^, which is more than two times the previous rate. From 2015 to 2019, the total biomass in Utah and Nevada remained largely unchanged, with a relatively slight decrease of 2,217,783 ± 37,535 Mg (2.218 ± 0.038 Mt), which is about 0.41% of the total biomass (Fig. S3 and Fig. 2a). Of this decrease, 1,721,553 ± 33,039 Mg (1.722 ± 0.022 Mt) was due to wildfires, accounting for roughly 78% of the overall reduction.

**Fig. 2.**
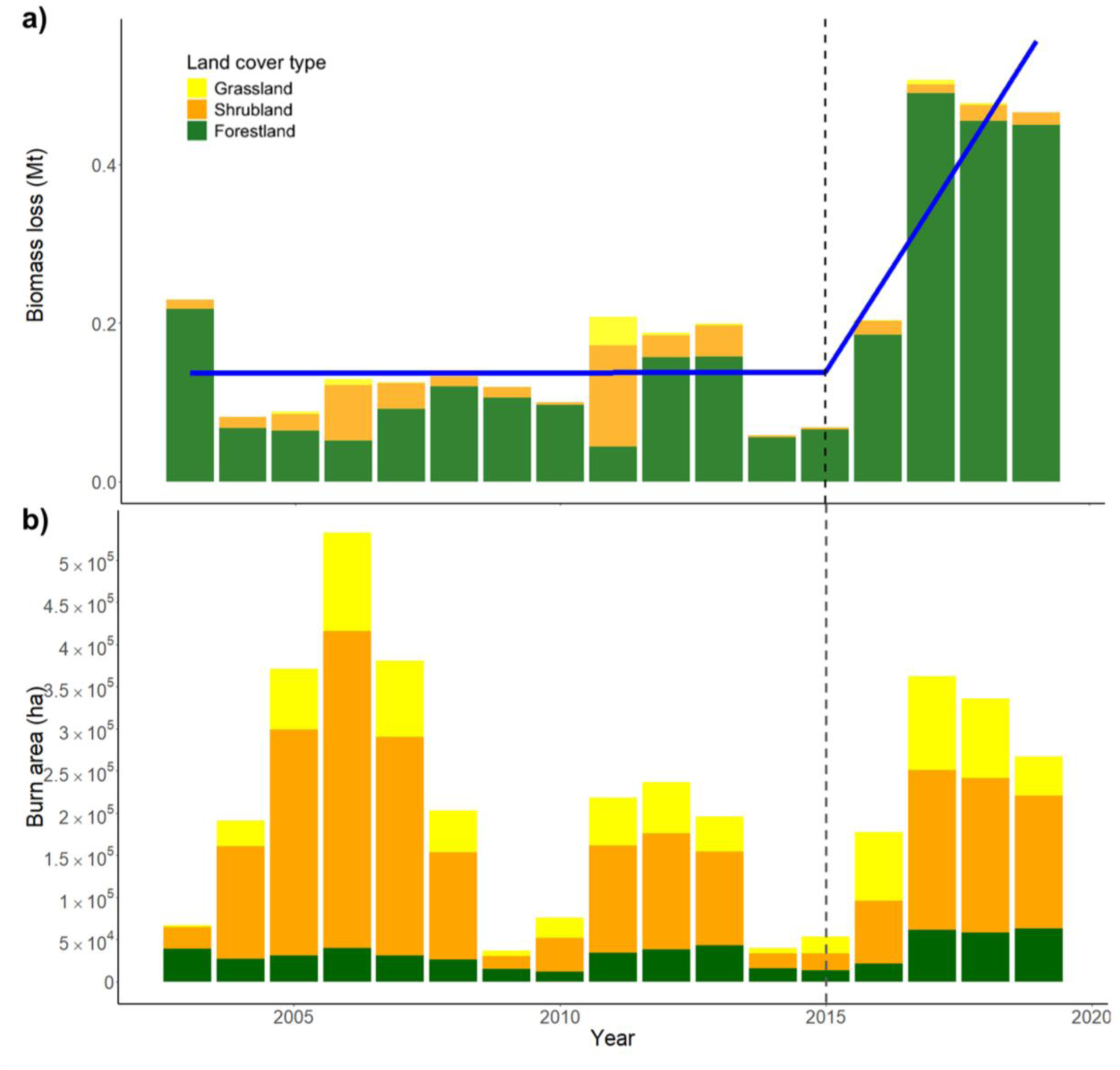
Trends in wildfire-induced biomass loss reveal a significant increase after 2015, but this trend is not reflected in the burn area. (a) Biomass loss caused by wildfires each year, which is the measurement we used in this study to assess the impact of wildfires. The height of each bar represents the total biomass loss (Mt), with different colors indicating the biomass loss for each land cover type separately. The dashed vertical line marks the tipping point by segmented regression, estimated to be in 2015, beyond which the biomass loss due to wildfires increases significantly. The blue lines represent the trend of biomass loss over the years. For the years before 2015, the biomass burnt by wildfires did not significantly change with the year (*β_1_* = 0 Mt yr^-1^, *p* > 0.1). In contrast, for the years after 2015, the biomass burnt by wildfires every year was positively and significantly correlated with the year (*β_1_*+*β_2_* = 0.105 Mt yr^-1^, *p* < 0.01). (b) Burn area of wildfires each year, which is the conventional measure that previous studies used to reflect the impact of wildfires. The height of each bar represents the total burn area (ha), with different colors indicating the burn area for each land cover type separately.

Forestland biomass loss was the primary contributor to total biomass loss from wildfires, accounting for an average of 82% and constituting more than half of the total biomass loss in most years (Fig. 2a). Furthermore, the segmented regression analysis shows that before 2015, the average forestland biomass loss was 102,470 ± 29,604 Mg yr^-1^; in contrast, after 2015, it increased by three fold, to 328,838 ± 31,590 Mg yr^-1^. In addition, apart from the net forestland biomass loss, there was a significant increase in the relative percentage of forestland loss after 2015, rising from an average of 76% over the previous 12 years to 94% thereafter. Additionally, before 2015, the biomass loss was 4,366 ± 2,114 Mg yr^-1^ in grassland and 31,239 ± 11,672 Mg yr^-1^ in shrubland; while after 2015, these values decreased to 1,939 ± 249 Mg yr^-1^ for grassland and 13,532 ± 1,390 Mg yr^-1^ for shrubland. Compared to the annual biomass loss from grassland and shrubland due to wildfires, the loss in forestland areas stands out as even more prominent which is completely different if analyzed by burn area (Fig. 2b). In most years, the burn area of woodland accounts for less than 50%, with an average contribution of only about 20.8% to the total burn area. In contrast, grassland contributes approximately 24.0%, while shrubland dominates with an average share of about 55.2% (Fig. S4).

From 2003 to 2019 (Fig. 2b), based on three-year moving average data, burn areas for shrubland, grassland, and forestland show that the year with the highest total burn area across all land cover types was 2006, with a combined area of 532,520 ha. In contrast, 2009 had the smallest total burn area, amounting to 36,614 ha. For shrubland, the largest burn area was in 2006, reaching 375,762 ha, while the smallest occurred in 2009 at 15,848 ha; For grassland, the maximum burn area was also in 2006, with 116,755 ha, and its minimum was in 2003 at 2,769 ha; For forestland, the highest burn area in 2019 at 62,882 ha, with the lowest recorded in 2010 at 11,735 ha. On average, the annual burn area was 133,030 ha for shrubland, 53,757 for grassland, and 33,492 ha for forestland, leading to an overall mean total burn area of 220,280 ha across all land cover types. The overall burn area displays a 6-year interannual cycle without an evident tipping point, while the burn area in forestlands has shown an upward trend since 2015.

### 3.3. Impacts of land cover types on wildfire biomass loss

Forestland fires, which naturally have higher biomass density before wildfires, tended to show greater severity, in terms of the amount of biomass density loss (Mg ha⁻¹) after wildfires. Wildfires with higher biomass density before the fire experienced greater biomass density loss, indicating more severe wildfire damage (Fig. 3a). Furthermore, areas with a higher percentage of forestland area typically had greater biomass density before a fire, resulting in more severe wildfire damage. These regions were also more likely to lose a significant amount of biomass and release more carbon if a wildfire occurs, making them especially vulnerable.

**Fig. 3.**
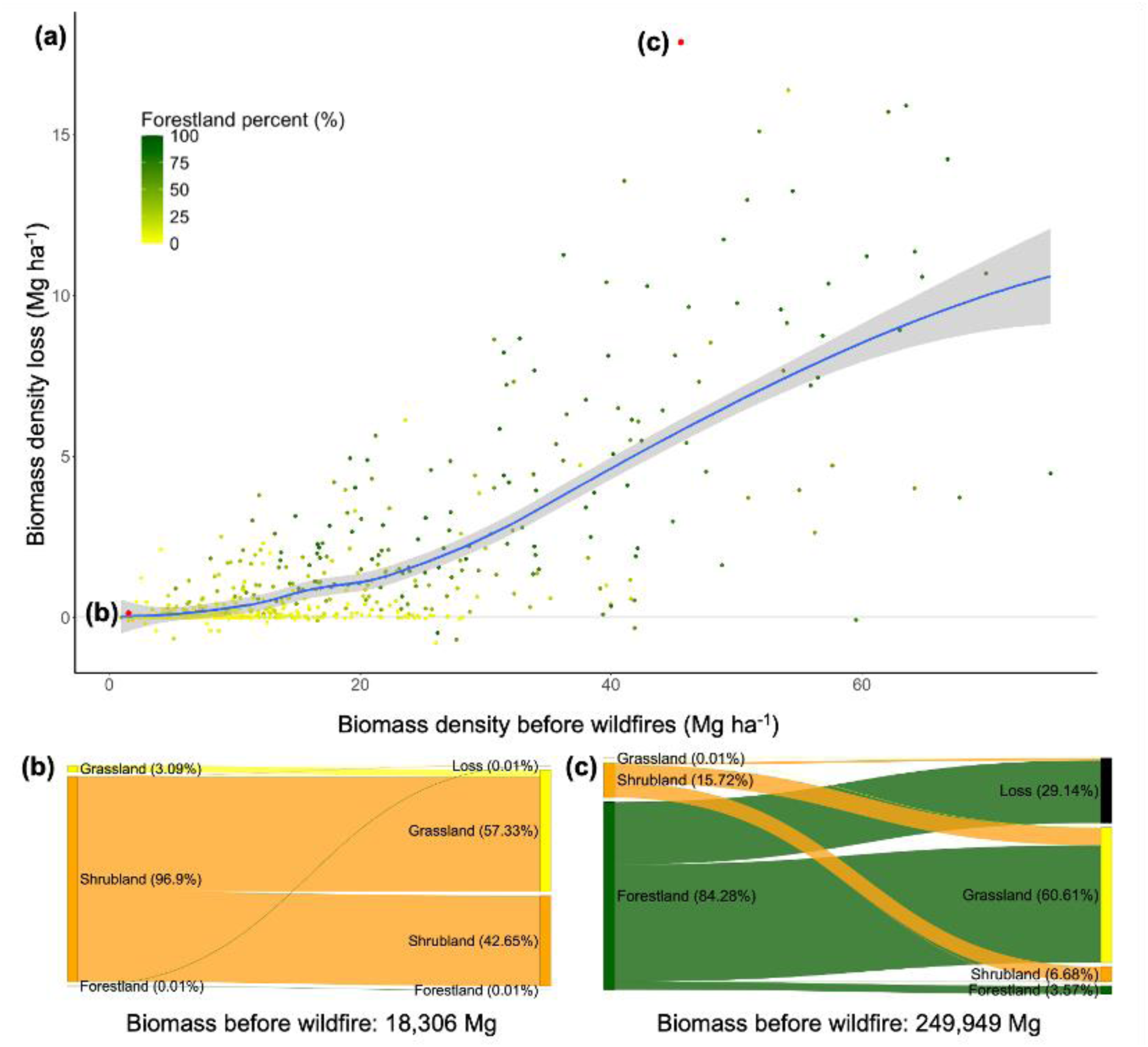
Relationship between land cover and wildfire-induced biomass loss. (a) Relationship between biomass density (Mg ha^-1^) and biomass density loss after wildfires (Mg ha^-1^), which is a quantitative measure of wildfire severity. Each point represents a single wildfire, with the color indicating the percentage of forestland area within the total burnt area. The impacts of land cover on wildfire-induced biomass loss are illustrated by two example wildfires: the Coyote fire in an area with a low percentage of forestland (b) and the West Valley fire in an area with a high percentage of forestland (c). Panels (b) and (c) are Sankey diagrams showing the transformation pathways of biomass from different land cover types before the wildfires (left side) to their states after the wildfires (right side).

We illustrate how wildfires burn biomass and release stored carbon across different land cover types with two examples from our analysis: the Coyote fire (Fig. 3b) and the West Valley fire (Fig. 3c). The former represents an area with lower forestland coverage, while the latter represents an area with higher forestland coverage. We found that the Coyote fire, which started on June 22nd, 2005 in Clark County, Nevada (36.500 °N, 114.994 °W) burned a total of 4,028 ha (Fig. 3b). The initial biomass was 18,306 Mg, primarily stored in shrubland areas. After the fire, 55.97% of the shrubland (10,245 Mg), which accounted for 54.23% of the total biomass, was converted into grassland. The wildfire resulted in a loss of only 0.01% (1.83 Mg) of the total biomass. Of this biomass loss, grassland contributed 5.97% (0.109 Mg), shrubland contributed 94.02% (1.72 Mg), and forestland contributed 0.01% (0.0002 Mg) of the total biomass loss. In contrast, the West Valley fire, which started on June 27th, 2018 in Washington County, Utah (37.441 °N, 113.387 °W), burned a total of 4,821 ha (Fig. 3c). The initial biomass was considerably higher at 249,949 Mg, with most of it stored in forestland areas. After the fire, 62.37% of the forestland (131,373 Mg), which accounted for 52.56% of the total biomass, was converted into grassland, and 29.14% (72,835 Mg) of the total biomass was lost due to the wildfire. Among this loss, grassland contributed 0.001% (0.728 Mg), shrubland contributed 3.44% (2,505 Mg), and forestland contributed 96.56% (70,329 Mg) of the total biomass loss. The contrasting examples of the Coyote and the West Valley fires showed how areas with higher forestland coverage tended to experience more severe biomass loss. Overall, the analysis shows that wildfire severity is closely linked to biomass density and land cover types, indicating that regions with denser biomass, especially forestlands, are more likely to experience substantial biomass loss during wildfires.

### 3.4 Impact of regional aridification on wildfire biomass loss across land cover types

Forestland ecosystems are uniquely sensitive to drought conditions, as evidenced by strong direct correlations between specific drought indices and biomass loss due to wildfires. By accounting for the interdependencies among 21 drought metrics, the MIIC algorithm effectively disentangled complex relationships and filtered out indirect associations, allowing for the identification of drought indices that exhibit direct correlations with biomass loss. The analysis revealed that such direct correlations occurred exclusively in forestland areas, while no statistically significant relationships were observed in shrubland or grassland regions. This finding underscores the distinct sensitivity of forestlands to drought-driven wildfire dynamics. Specifically, six drought indices—PDSI, 1-year SPEI, 1-year SPI, 180-day EDDI, 90-day SPEI, and 90-day SPI—were found to be directly correlated with biomass burned in forestlands. By isolating these key indicators from a highly interdependent set of variables, the MIIC algorithm enabled a more precise understanding of the role of drought in influencing wildfire behavior in forested landscapes.

To further understand how aridification prior to wildfires differs across land cover types, we analyzed the temporal trends of MIIC-identified drought indices, which showed a clear trend of increasing aridification in forestland ecosystems. The interannual analysis shows that in grassland and shrubland areas, these drought indices exhibited no clear drying or wetting trends before wildfires (Fig. 4). In contrast, forestland regions exhibit a consistent negative trend in PDSI, 90-day SPEI, and 90-day SPI, indicating increasing dryness leading up to wildfire events. This pattern of pre-fire aridification appears to be specific to forestland ecosystems, underscoring their heightened sensitivity to drought conditions. Notably, similar negative trends were also observed in additional drought indices not identified as having direct associations by MIIC, suggesting potential secondary relevance (Fig. S5).

**Fig. 4.**
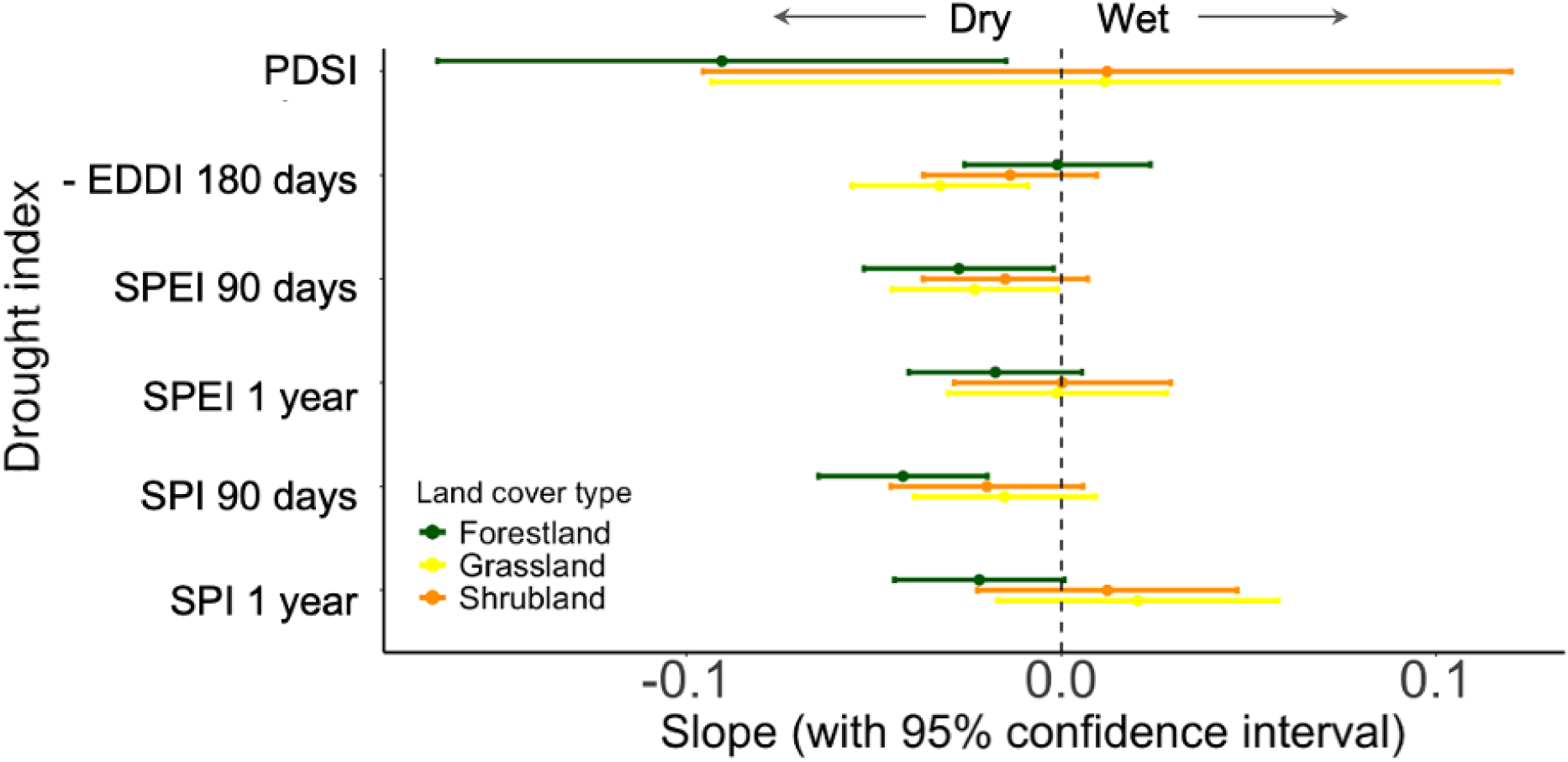
Varying aridification levels across different land cover types. The y-axis shows the slope of the linear trend of drought indices over the years from 2003 to 2019, representing the rate of change in dryness conditions across time. More negative slopes indicate increasing aridification. The interannual variability of four drought indices (PDSI, SPEI, SPI, and negative EDDI) is shown across land cover types, along with their 95% confidence intervals. For all indices, lower values denote drier conditions..

## 4. Discussion

We found that biomass loss in Utah and Nevada due to wildfires has significantly increased over the past two decades. This trend is largely attributed to more intense wildfires, driven by climate-induced aridification and prolonged droughts. An increasingly dry climate emerges as the primary driver of rising wildfire biomass loss by increasing the incidence of wildfires in relatively biomass-dense forestlands, as evidenced through the dependency detection of drought and wildfire data. We also find that land cover plays an important role in wildfire impacts, with forests experiencing greater biomass losses compared to other areas. Notably, the year 2015 serves as a tipping point; after 2015, annual wildfire biomass loss increased significantly. Rising forestland wildfire biomass loss further contributed to the growing biomass loss during wildfires. The time of wildfire-induced biomass loss is gradually shifting later into the year (SI §11), signaling that late-season, high-biomass-loss forestland wildfires demand extra attention, while the non-forestland wildfires do not have a significant change. Overall, our findings provide valuable insights for evaluating wildfire carbon emissions by offering a quantitative analysis of biomass loss trends, complementary to burn area trends, and their driving factors. These insights have important policy implications, particularly for mitigating wildfire risks and adapting to changing climate conditions.

### 4.1. Rising biomass loss and the impact of forestland wildfires

Supporting our first hypothesis, we found that, over the past two decades, biomass loss due to wildfires in Utah and Nevada has significantly increased, with forestlands emerging as the primary contributors to this trend. Particularly after 2015, which marked a tipping point, the overall biomass lost due to wildfires began to increase rapidly. Notably, this increase in biomass loss does not coincide with a similar breakpoint in burned area per year, suggesting that there is a significant change in the nature of wildfire in this region around 2015.

Since 2015, the increasing proportion of biomass loss from forestlands has emerged as a key driver of the overall upward trend. Our results underscore that burned area alone provides an incomplete picture of wildfire impact, as it does not account for differences in biomass density across land cover types. In our study, we find that, although wildfires often burn larger areas in land covers such as grasslands or shrublands (Fig. 2), which is also demonstrated by other research (Zhai et al., 2023), their lower biomass densities result in comparatively smaller losses (Fig. 3). In contrast, forestlands, with their high biomass density, contribute disproportionately to total biomass loss when affected by wildfires (Fig. 2). This pattern is especially evident in the significant rise in burned forestland areas since 2015, which closely mirrors the observed increase in biomass loss (Fig. 2). Importantly, this trend cannot be explained by changes in total biomass, which has remained relatively stable over the same period (Fig. S3).

### 4.2. Impact of aridification on wildfire biomass loss

Consistent with the hypothesis that climate change–induced aridification leads to greater biomass loss during wildfires, our results show that fire-affected landscapes have become increasingly drier in recent years (Fig. 4), especially in forestland areas, where biomass losses have also intensified. This finding aligns with previous studies on wildfires in the western US (Abatzoglou & Williams, 2016; Goss et al., 2020; Williams et al., 2019) and Canada (Byrne et al., 2024). Our finding suggests that aridification has specifically intensified biomass loss in forestlands during wildfires, which aligns with the existing study (Wang et al., 2024), where warmer summers and drier winters significantly increased forest exposure, fire severity, and burned area in California; meanwhile, we found no significant impact of aridification on non-forestland areas. This aligns with our observation (Fig. S6) that only the burned area of forestlands has increased, while the burned area in non-forestland regions continues to follow a cyclical pattern. Additionally, based on our MIIC analysis, drought indices are related only to biomass loss in forestlands during wildfires and have no association with biomass loss in non-forestland areas. This variation in sensitivity to aridification due to differences in land cover has also been shown in California (Williams et al., 2019). The different sensitivities might depend on the varying ignition thresholds of different land cover types (Li et al., 2019; Newberry et al., 2020). Aridification increases fuel aridity (Abatzoglou & Williams, 2016), and thus significantly lowers the ignition threshold of forestlands. However, this increase in fuel aridity may have less impact on the ignition threshold of non-forestland areas, where fuel dries more easily and reaches its ignition threshold early in the fire season, than that of forestland areas. What’s more, the increase in biomass loss in forestland has also shifted the distribution of wildfire-induced biomass loss within a year, from earlier-season fires in low-biomass grasses and shrubs to later-season fires in higher-biomass forestlands (SI §11).

### 4.3. Caveats and future work

Our analysis provides one of the most comprehensive assessments of the impact of wildfires on aboveground biomass and land cover types across a large region. However,this study examines the immediate impact of wildfires on different land cover types, but it does not consider the prolonged effects of wildfires, such as secondary succession after a wildfire. Long-term impact is also important because prolonged effects can significantly alter the recovery process and final state of ecosystems after wildfires. For instance, secondary succession can lead to changes in species composition and ecosystem functions (B. Liu et al., 2024; Seidl & Turner, 2022), which may take decades or even longer to manifest (Coradini et al., 2022). Since forestlands have relatively long growth cycles compared to grasslands or shrublands, their recovery time after experiencing wildfires is also longer (Abbas et al., 2023), making the prolonged effects on forestlands particularly significant. Additionally, prolonged effects include changes in soil (Baur et al., 2024; Pellegrini et al., 2022), nutrient cycling (Pellegrini et al., 2015), and hydrological processes (Baur et al., 2024; Williams, Livneh, et al., 2022), which are crucial factors for ecosystem health and stability. Excluding these prolonged effects might lead to an underestimation of the long-term ecological impacts of wildfires, thereby affecting the development of management and conservation measures. Therefore, future research should focus more on the prolonged effects of wildfires to fully understand their ecological impacts on different land cover types.

Second, this study stands out for its high spatial resolution (30 meters) and broad geographic coverage in analyzing biomass loss and carbon emissions from U.S. wildfires, but it ends in 2020, lacking the most recent wildfires. As climate extremes have become more common, the study may not fully capture how wildfire severity, in terms of biomass loss, has changed under the most recent climate change. Recent events suggest that wildfire severity may have even increased in response to accelerating climate change (Wang et al., 2024). The unprecedented wildfires in Canada in 2023 (Byrne et al., 2024). and the severe fires in the Los Angeles metropolitan area in early 2025 (Swain et al., 2025) highlight this emerging trend. These cases underscore the importance of extending temporal coverage in future research to include the most recent years, which is essential for understanding how wildfire dynamics are shifting under current climate conditions.

Third, at the time this study was conducted, the annual NLCD maps were not yet available, and the corresponding validation results had not been published. Consequently, we adopted the most recent, well-established NLCD map from the year closest to the wildfire event as a proxy for pre-fire land cover conditions. While this approach may not fully capture subtle temporal changes in land cover, it provides a level of spatial and thematic accuracy sufficient for the objectives of this research. In future work, the incorporation of updated, annually resolved NLCD datasets will allow for a more precise characterization of land cover dynamics, thereby improving the assessment of wildfire impacts and enabling a more robust evaluation of temporal trends.

Finally, due to the lack of suitable ground-based measurements for grassland and shrubland ecosystems, it was not possible to directly validate the terraPulse aboveground biomass estimates for these land-cover types in this study. This limitation should be considered when interpreting the results for non-forest vegetation. Nevertheless, previous research by Cao et al. (2015) has demonstrated that terraPulse aboveground biomass estimates for forested areas are consistent with Forest Inventory and Analysis (FIA) data, indicating a high degree of reliability in forestland applications. This provides confidence in the methodology’s robustness, while also highlighting the need for more comprehensive ground observations in grassland and shrubland ecosystems to further improve cross-ecosystem validation.

Despite these limitations, our work yields policy-relevant insights. First, detecting biomass-loss tipping points reveals when ecosystems transition into persistently higher-severity fire regimes. These long-term shifts have direct implications for strategic land management: (i) scaling up prescribed fire and cultural burning across landscapes approaching critical thresholds, (ii) expanding multi-year mechanical thinning programs in high-risk zones, and (iii) adjusting budgets to prioritize regions showing increasing biomass loss trends. This is consistent with recent findings that proactive fuels management reduces subsequent fire severity and enhances operational opportunities (Davis et al., 2023, 2024; Wu et al., 2023). Second, within each year, the identification of high-impact “heat points” in biomass loss pinpoints weeks or months when the potential ecological impact of burning is greatest. These intra-annual windows can inform tactical measures: pre-positioning crews and equipment, timing prescribed burns to reduce fuel before high-impact periods, implementing smoke-management protocols, and enhancing public communication when ecosystem damage risk is elevated. By jointly leveraging tipping points and intra-annual heat points, agencies can integrate long-term resilience planning with short-term fire-season readiness, ultimately improving the efficiency and effectiveness of wildfire management.

### 4.4. Conclusions

This study underscores the growing threat of wildfire-related biomass loss in the western US. Our study highlights a significant increase in biomass loss due to wildfires in Utah and Nevada, with a critical tipping point observed in 2015. The finding reveals a novel shift in the western US fire regime, where wildfire biomass loss began to intensify notably. This marked the beginning of a sharp rise in wildfire-related biomass loss, predominantly driven by forestland areas. Our findings underscore that these forestlands have consistently been the primary contributors to the overall biomass loss during wildfires, with climate change, especially aridification, becoming the main driver of this trend. Considering that Utah and Nevada are rapidly growing population areas and experience severe wildfires, similar to other regions in the western US, the findings of this study can inform wildfire management and policy strategies. By integrating these insights, stakeholders can better anticipate wildfire risks, with the potential for a prolonged wildfire season and increased forestland biomass loss.

