## Supplemental Materials for "Accelerated biomass loss in western US forestlands due to shifting fire regime"

#### 1. Quantify the overall uncertainty in biomass estimates

We used Monte Carlo simulations to quantify the overall uncertainty in biomass estimates for specific regions in Utah and Nevada. We conducted the MCS using the following steps: Each pixel $i$ within the study area from the terraPulse dataset has a biomass value $B_i$ and its corresponding uncertainty $SD_i$, expressed as the standard deviation (SD). To perform the simulations, we generated random values $b_i^k$ for the biomass of each pixel, drawing from a distribution defined by the pixel's mean biomass value and its SD:

$$b_i^k \sim N(B_i, SD_i)$$

Then we computed the total biomass for the iteration $k$ as:

$$T_k = \sum_{i=1}^{N} b_i^k$$

Where $k$ ranges from 1 to $M$, and $M$ is the total number of simulations. $N$ is the total number of pixels. This process was repeated for each pixel to simulate a total biomass value for the entire study area. We conducted a total of 2000 simulations, each producing a distinct total biomass estimate for the regions of interest. From these simulations, we calculated the mean total biomass:

$$T_{mean} = \frac{1}{M} \sum_{k=1}^{M} T_k$$

and the associated uncertainty of the overall biomass:

$$SD_T = \sqrt{\frac{1}{M-1}\sum_{k=1}^{M}(T_k - T_{mean})^2}$$

This method provides a robust statistical approach to understanding the variability and confidence levels in biomass assessments.

#### 2. Drought index

| Drought index | Defination |
|---|---|
| Standardized precipitation index (SPI)(Climate data guide, 2025) | $SPI = \frac{(P-P^*)}{\sigma_P}$<br>where $P$ = precipitation<br>$P^*$ = mean precipitation<br>$\sigma_P$ = standard deviation of precipitation |
| Evaporative drought demand index (EDDI)(Physical sciences laboratory, 2025) | The Evaporative Demand Drought Index (EDDI) is an experimental drought monitoring and early warning guidance tool. It examines how anomalous the atmospheric evaporative demand is for a given location and across a time period of interest. |
| Standardized precipitation evapotranspiration index (SPEI)(Climate data guide, 2025) | The procedure for calculating the SPEI is similar to that for the SPI. However, the SPEI uses the difference between precipitation and reference evapotranspiration ($P - ETo$), rather than precipitation (P) as the input. |

| | |
|---|---|
| Palmer drought severity index (PDSI)(Climate data guide, 2025) | The PDSI has been reasonably successful at quantifying long-term drought. As it uses temperature data and a physical water balance model, it can capture the basic effect of global warming on drought through changes in potential evapotranspiration. Monthly PDSI values do not capture droughts on time scales less than about 12 months. |
| Palmer Z Index (Z)(Physical sciences laboratory, 2025) | The Palmer Z Index measures short-term drought on a monthly scale. |

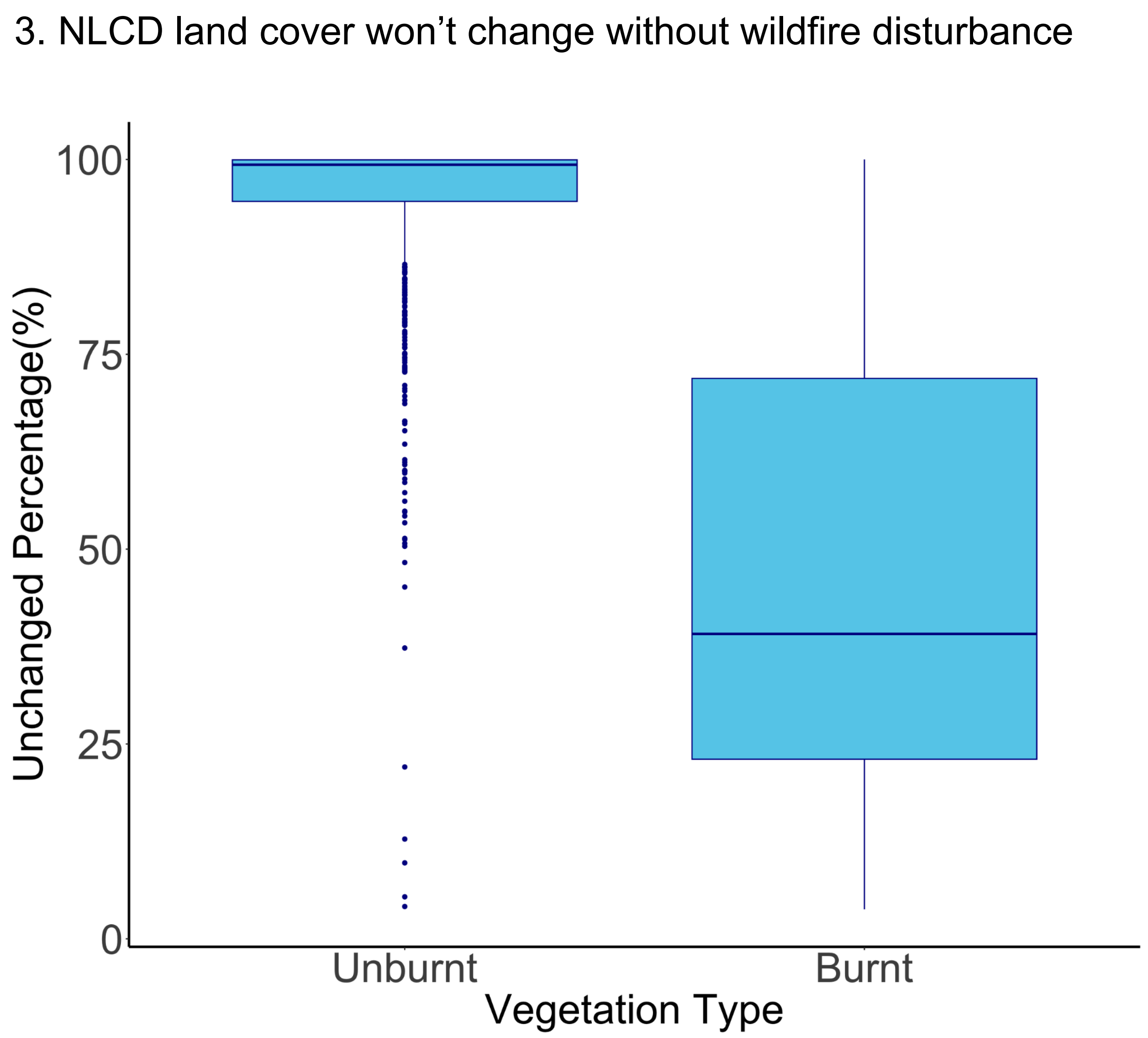


**Fig. S1 | Land cover changes for places with and without fires.** The two box plots show changes in land cover under wildfire and non-wildfire conditions. The burned polygons are derived from the MTBS dataset. The unburned areas are selected from within the bounding boxes of the wildfire polygons but outside the actual burned areas. These unburned regions are assumed to have similar environmental conditions to the corresponding burned areas and are used as control plots.

### 4. Examples from terraPulse biomass product

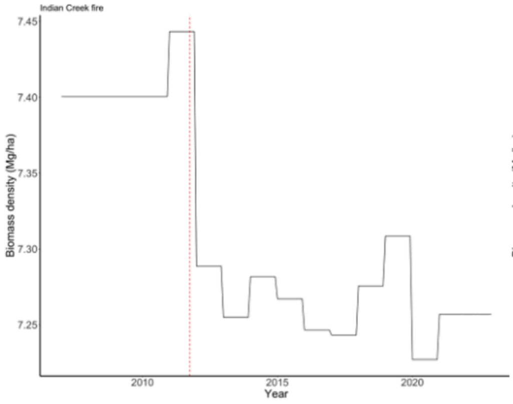


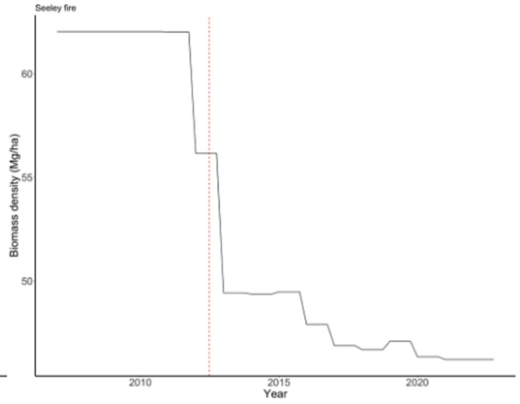


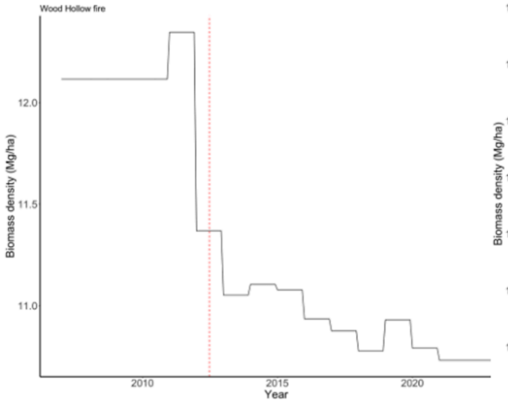


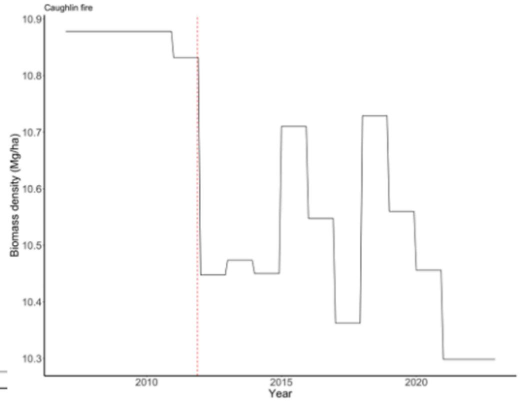


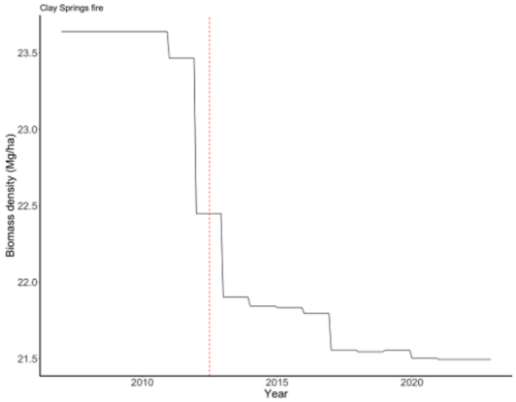


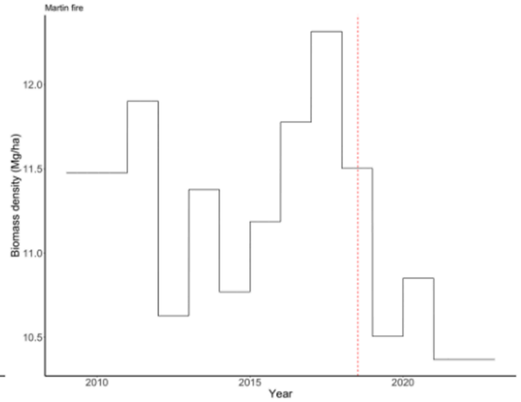


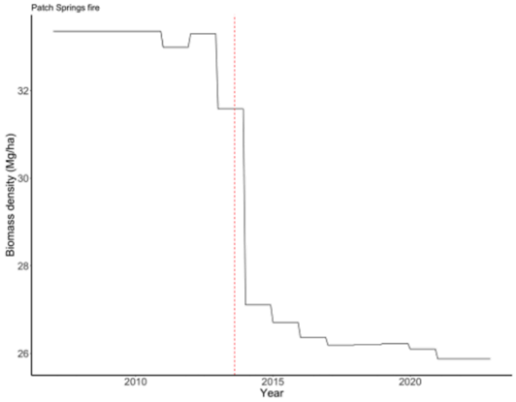


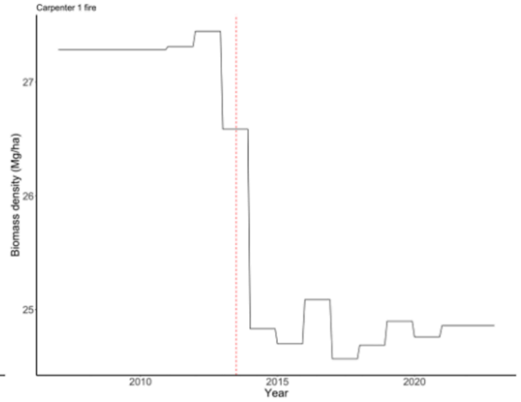

**Fig. S2 | Biomass density change under wildfire disturbance.** The 8 panels show the annual biomass density (Mg ha$^{-1}$). The dashed red line represents the time of wildfire disturbance.

#### 5. Segmented regression

Firstly, the equation for a segmented regression pinnacle can be expressed as :

$$\mu = E[Y] = \beta_1 z + \beta_2 (z - \psi)_+$$

Where $(z - \psi)_+ = (z - \psi) \times I(z > \psi)$ and $I(\cdot)$ is the indicator function, which is 1 if $z > \psi$ and 0 otherwise. $\mu = E[Y]$ is the expected value (mean) of the response variable $Y$. $\beta_1$ is the slope before the breakpoint. $\beta_2$ is the change in the slope after the breakpoint. $\psi$ is the breakpoint. Then we made an initial guess for the breakpoint $\overline{\psi}$ (we guessed 2014) and used iterative algorithms to estimate the model parameters. In each iteration, we fit the following linear model (Muggeo 2003; 2008):

$$\mu = \beta_1 z + \beta_2 (z - \overline{\psi})_+ + \gamma I(z > \overline{\psi})^-$$

Where $I(\cdot)^- = - I(\cdot)$ and $\gamma$ is the parameter that may be understood as a re-parameterization of $\psi$ and therefore accounts for the breakpoint estimation. During each iteration, this standard linear model is fitted, and the value of the breakpoint is updated by:

$$\widehat{\psi} = \overline{\psi} + \widehat{\gamma} / \widehat{\beta_2}$$

where $\overline{\psi}$ is the unadjusted breakpoint estimate, $\widehat{\gamma}$ is the breakpoint adjustment term, and $\widehat{\beta_2}$ is the current estimate of the change in slope. When $\widehat{\gamma}$ is small, it means that the algorithm has converged and we can get the tipping point $\widehat{\psi}$. Finally, we employed the Davies (Davies 1987) test to determine the existence of a breakpoint and obtain the corresponding p-value.

#### 6. Weighted quantile regression

By optimizing the following equation, we obtained the regression parameters (a vector consisting of slope and intercept):

$$\beta_\tau = \underset{\beta_\tau}{argmin} \left[- \sum_{y_i < x_i^T\beta} w_i(1 - \tau)(y_i - x_i^T\beta) + \sum_{y_i \geq x_i^T\beta} w_i\tau(y_i - x_i^T\beta)\right]$$

Where β is the vector of regression parameters. $y_i$ is the response variable for the $i^{th}$ observation. $x_i$ is the vector of the explanatory variable for $i^{th}$ observation. τ is the quantile selected. $w_i$ is the weight for $i^{th}$ observation. In this study, $y$ represents the week when wildfires occurred, $x$ represents the year when wildfires occurred, and $w$ represents the biomass burned by the wildfires. We used $\tau = 0.05$ to mark the beginning of the period of wildfire-induced biomass loss (5% quantile) and $\tau = 0.95$ to mark its end (95% quantile).

#### 7. Changes in total aboveground live biomass in Utah and Nevada over nineteen years.

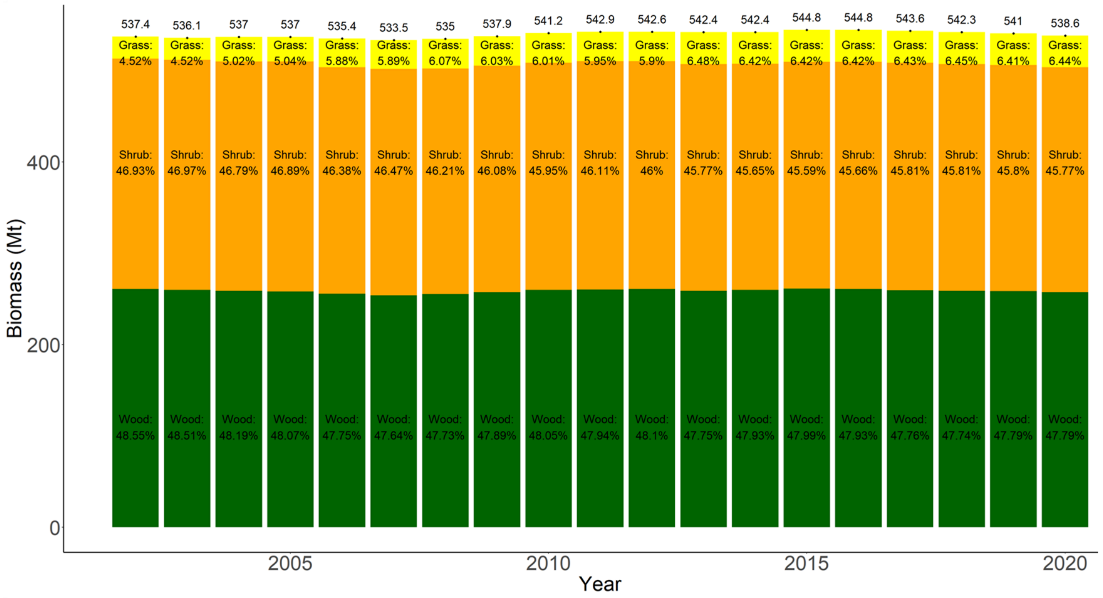


**Fig. S3 | Changes in total aboveground live biomass in Utah and Nevada.** The stacked columns represent the aboveground live biomass for different land covers: forestland, Shrubland, and Grassland. Each stack column representing different land cover for each year has a segment proportion that indicates the percentage of that biomass contributing to the total biomass. The number at the top of each stack column represents the total weight of biomass in the study area for that year, measured in Mt.

### 8. Wildfire-induced biomass loss and burn area of different land cover types.

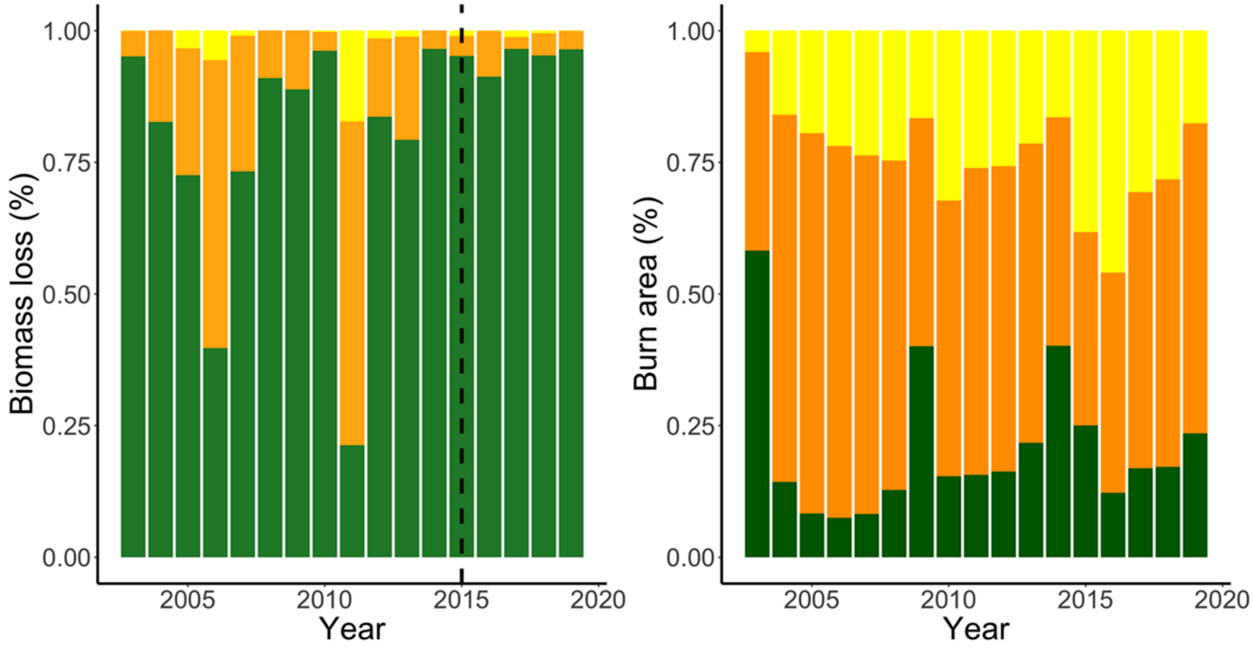


**Fig. S4 | Annual biomass loss and burn area percentage of different land cover types.** The stacked columns on the left represent the percentage of aboveground live biomass loss in each year's wildfires for different land covers: Forestland, Shrubland, and Grassland. The stacked columns on the right represent the percentage of burn area in each year's wildfires for different land covers.

#### 9. Drought indices

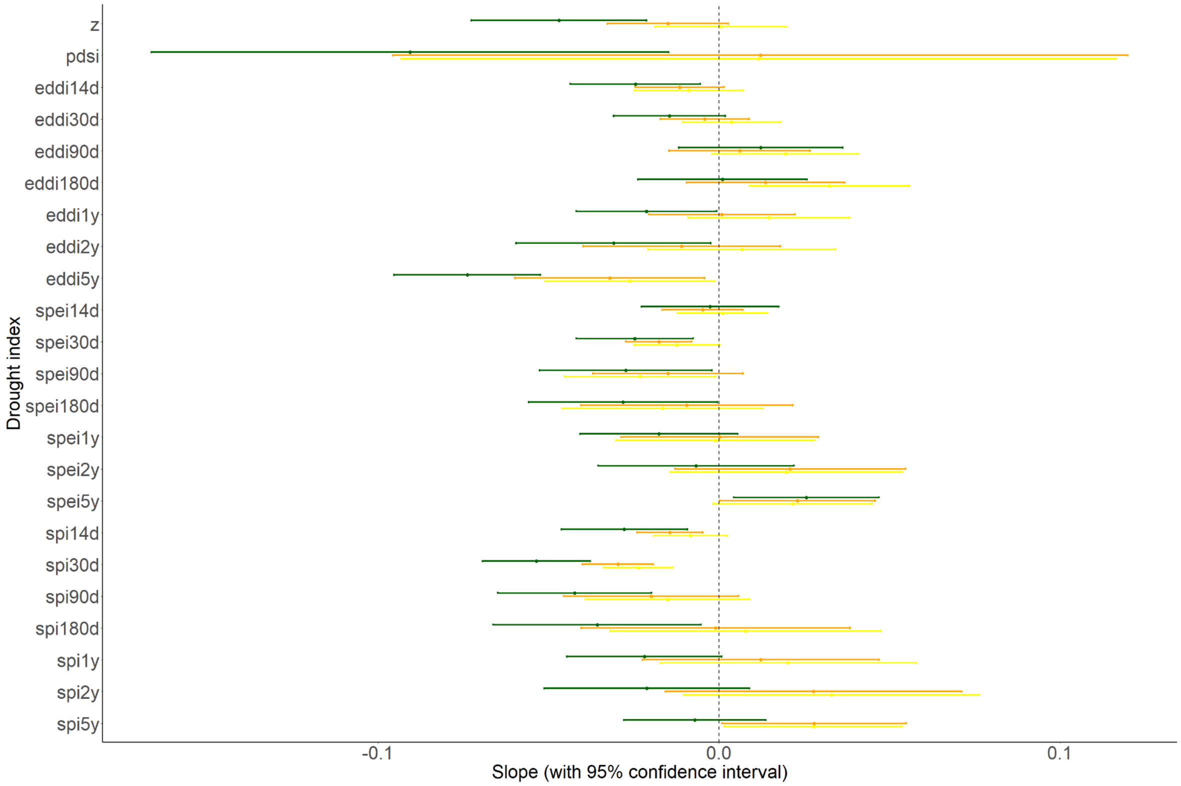


**Fig. S5 | Varying (all) aridification levels across different land cover types.** The y-axis shows the slope of the linear trend of drought indices over years, representing the rate of change in dryness conditions across time. More negative slopes indicate increasing aridification. The interannual variability of four drought indices (PDSI, SPEI, SPI, and negative EDDI) is shown across land cover types, along with their 95% confidence intervals. For all indices, lower values denote drier conditions..

10. Increasing area of forestland burned by wildfires

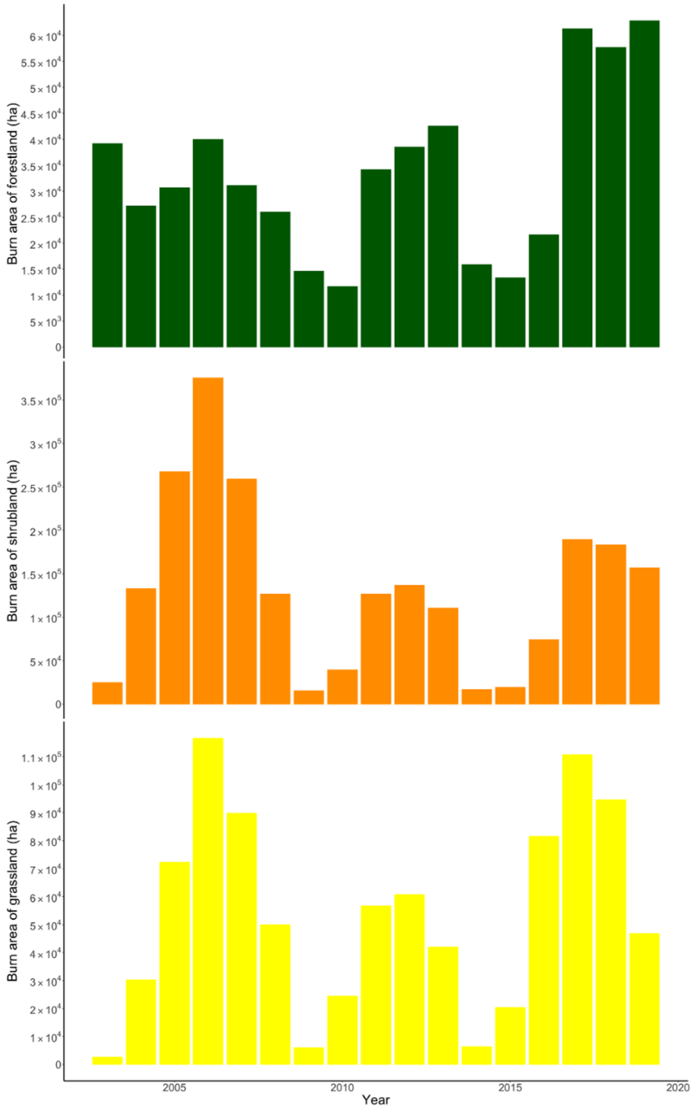


**Fig. S6 | Wildfire burn area in Utah and Nevada.** (a)The area burnt over each year. The height of each column represents the total area burnt (ha), with different colors indicating the area burnt for each land cover type separately. (b)The annual trends in the area burned in forestland areas.

### 11. Timing of biomass loss attributed to wildfires

#### 11.1. Temporal and impact-based quantification of wildfire biomass loss

In this study, the heat point of wildfire-induced biomass loss (HP-WBL) was determined by both the timing of wildfires and the biomass loss they caused. The ignition of a wildfire is a stochastic event; our exploratory data analysis also suggested that wildfires can occur at any time of the year. Since the timing of the first and last wildfires may not correspond to detectable biomass loss, using these dates to define the HP-WBL may not accurately capture the underlying patterns and processes of biomass change. To determine the HP-WBL for each year, we used quantile regression weighted by the biomass loss of each wildfire. Specifically, we identified the 5th percentile of total biomass loss as the start of the HP-WBL and the 95th percentile as the end. This approach defined the wildfire HP-WBL as the period during which 90% of the biomass loss occurred. By weighting the regression with biomass loss, larger and more destructive fires had a greater influence on determining the HP-WBL. This approach emphasized severe wildfires, highlighting the periods with the most substantial ecological and economic consequences.

We conducted the weighted quantile regression (SI §6) in the following form:

$$Q_\tau(y_i \mid x_i) = \beta_0^{(\tau)} + \beta_1^{(\tau)} x_i$$

where $Q_\tau(y_i \mid x_i)$ represents the $\tau - th$quantile of the distribution of $y_i$given $x_i$, $\beta_0^{(\tau)}$ is the intercept for the $\tau - th$ quantile, $\beta_1^{(\tau)}$ is the slope for the $\tau - th$ quantile. By specifying (τ =

0.05), we estimate the model for the 5th quantile. Similarly, by specifying (τ = 0.95), we estimate the model for the 95th quantile.

#### 11.2. Temporal shift of wildfire-induced biomass loss

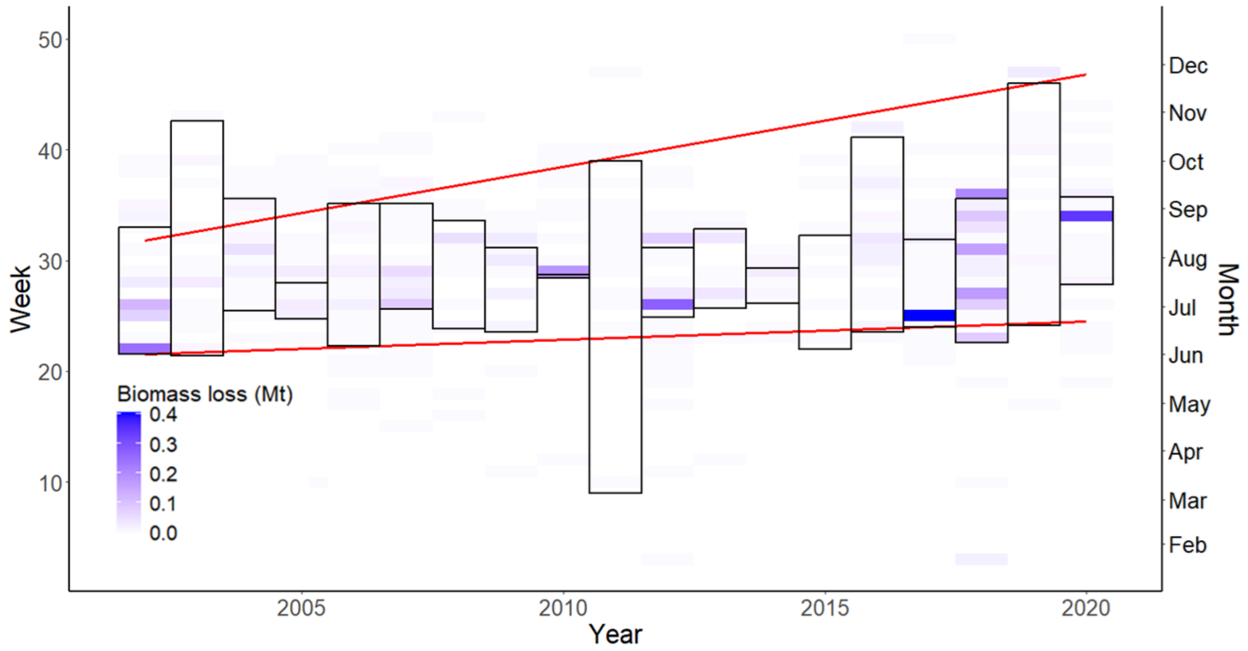


**Fig. S7 | Annual HP-WBL dates weighted by biomass loss.** Each vertical box marks the duration of the HP-WBL, defined as the period during which 90% of the biomass loss caused by wildfires has occurred (5% before and 5% after). The heat map colors represent biomass loss for each week of each year. The two red lines show the results of quantile regressions at the 5th and 95th percentiles, weighted by biomass loss. The start of the HP-WBL is defined by the 5th percentile, and the end is defined by the 95th percentile. The horizontal axis represents the years corresponding to the columns in the figure, the left vertical axis represents the week of the year, and the right vertical axis represents the month of the year.

Each year's HP-WBL shows notable changes in timing and duration. Based on the HP-WBL dates identified through weighted quantile regression, it is evident that the start and end dates of each HP-WBL, represented by the black regions in Fig. S6, occurred within the same calendar year. In other words, our study area did not experience HP-WBLs overlapping multiple years. The quantile regression of HP-WBL (Fig. S7) reveals a lengthening of the duration of HP-WBL over the years. Specifically, the coefficient ($\beta_1^{(0.05)}$) for the start of the HP-WBL ($\tau = 0.05$) is 0.162 ± 0.107 wk $yr^{-1}$ ( $p > 0.1$), indicating there was no significant delaying or advancing of the starting date of HP-WBL. Conversely, the coefficient ($\beta_1^{(0.95)}$) for the end of the HP-WBL ($\tau = 0.95$), is 0.835 ± 0.359 wk $yr^{-1}$ ($p < 0.05$), indicating a significant annual delay of approximately 0.835 weeks.  The HP-WBL in 2002 began at the very start of June and ended at the beginning of August. In contrast, the HP-WBL in 2020 started at the end of June and ended at the end of November. Between 2002 and 2020, the start of the HP-WBL showed no significant change, and the end was delayed by 15.03 ± 6.46 weeks, indicating the prolonging of the HP-WBL by 15.03 ± 6.46 weeks.

Climate change will not only increase biomass loss, but, as our findings suggest, also prolong HP-WBL. We find a clear pattern of the end of HP-WBL occurring progressively later in the year and no significant change in the start of HP-WBL, likely influenced by climate change (Ellis et al. 2022; Jolly et al. 2015), which results in lower fuel moisture levels (Halofsky et al. 2020). Notably, the larger slope for the end of HP-WBL compared to the start shows an extended HP-WBL in recent years.

Wildfires have predominantly affected shrubland and grassland areas, which have lower ignition thresholds compared to forestland areas (Li et al. 2019; Newberry et al. 2020). As a result, these types of fires usually mark the start of HP-WBL. Our recent observations also indicate that the expanding burning in forestland areas is contributing to a lengthening of HP-WBL (Fig. S6). Forestland, having a higher ignition threshold, often catches fire later, after the accumulation of dry, combustible biomass (Croker et al. 2023). Additionally, forestland fires are associated with significant biomass loss, so when wildfire timing is weighted by biomass loss, the end of HP-WBL shifts significantly later. In contrast, since biomass loss from grassland and shrubland wildfires remains relatively stable, the start of HP-WBL has no significant change.
